# shinyheatmap: ultra fast low memory heatmap web interface for big data genomics

**DOI:** 10.1101/076463

**Authors:** Bohdan B. Khomtchouk, James R. Hennessy, Claes Wahlestedt

**Affiliations:** Center for Therapeutic Innovation and Department of Psychiatry and Behavioral Sciences, University of Miami Miller School of Medicine, 1120 NW 14th St., Miami, FL, USA 33136; Department of Mathematics, University of Miami, 1365 Memorial Drive, Coral Gables, FL, USA 33146

## Abstract

**Background:** Transcriptomics, metabolomics, metagenomics, and other various next-generation sequencing (-omics) fields are known for their production of large datasets, especially across single-cell sequencing studies. Visualizing such big data has posed technical challenges in biology, both in terms of available computational resources as well as programming acumen. Since heatmaps are used to depict high-dimensional numerical data as a colored grid of cells, efficiency and speed have often proven to be critical considerations in the process of successfully converting data into graphics. For example, rendering interactive heatmaps from large input datasets (e.g., 100k+ rows) has been computationally infeasible on both desktop computers and web browsers. In addition to memory requirements, programming skills and knowledge have frequently been barriers-to-entry for creating highly customizable heatmaps.

**Results:** We propose shinyheatmap: an advanced user-friendly heatmap software suite capable of efficiently creating highly customizable static and interactive biological heatmaps in a web browser. shinyheatmap is a low memory footprint program, making it particularly well-suited for the interactive visualization of extremely large datasets that cannot typically be computed in-memory due to size restrictions. Also, shinyheatmap features a built-in high performance web plug-in, fastheatmap, for rapidly plotting interactive heatmaps of datasets as large as 10^5^ − 10^7^ rows within seconds, effectively shattering previous performance benchmarks of heatmap rendering speed.

**Conclusions:** shinyheatmap is hosted online as a freely available web server with an intuitive graphical user interface: http://shinyheatmap.com. The methods are implemented in R, and are available as part of the shinyheatmap project at: https://github.com/Bohdan-Khomtchouk/shinyheatmap. Users can access fastheatmap directly from within the shinyheatmap web interface, and all source code has been made publicly available on Github: https://github.com/Bohdan-Khomtchouk/fastheatmap.

## Introduction

Heatmap software can be generally classified into two categories: static heatmap software [30, 29, 49, 25, 14, 7, 8, 16, 19] and interactive heatmap software [32, 4, 18, 51, 23, 34, 48, 1, 17, 9, 52]. Static heatmaps are pictorially frozen snapshots of genomic activity displayed as colored images generated from the underlying data. Interactive heatmaps are dynamic palettes that allow users to zoom in and out of the contents of a heatmap to investigate a specific region, cluster, or even single gene while, at the same time, being able to hover the mouse pointer over any specific row and column entry in order to glean information about an individual cell’s contents (e.g., gene name, expression level, and column name). Interactive heatmaps are especially important for visualizing large gene expression datasets wherein individual gene labels eventually become unreadable due to text overlap, a common drawback seen in static heatmaps of large input data matrices. As such, interactive heatmaps are popular for examining the entire landscape of a large gene expression dataset while, at the same time, allowing users to zoom into specific sectors of the heatmap to visualize them in a magnified manner (i.e., at various resolution levels). Currently, there is a pressing need for modern libraries that are able to visually scale millions of data points at various resolutions [43]. In general, new software infrastructure that facilitates interactive navigation and smooth scaling at different resolution levels is necessary for on-the-fly calculations of both the frontend and backend algorithms in big data visualization software [42].

Even though static heatmaps are still the preferred type of publication figure in many studies, interactive heatmaps are becoming increasingly adopted by the scientific community to emphasize and visualize specific sectors of a dataset, where individual numerical values are rendered as user-specified colors. As a whole, the concept of interactivity is gradually shifting the heatmap visualization field into data analytics territory, for example, by synergizing interactive heatmap software with integrated statistical and genomic analysis suites such as PCA, differential expression, gene ontology, and network analysis [22, 17]. However, currently existing interactive heatmap software are limited by implicit restrictions on file input size, which functionally constrains their range of utility. For example, in Clustviz [22], which employs the pheatmap R package [19] for heatmap generation, input datasets larger than 1000 rows are discouraged [20] for performance reasons. Similarly, in MicroScope, the user is prompted to perform differential expression analysis on the input dataset first, thereby shrinking the number of rows rendered in the interactive heatmap to encompass only statistically significant genes [17]. In general, the standard way of thinking has been to avoid the production of big heatmaps due to a combination of various factors such as poor readability, as static heatmaps are not zoomable; computational infeasibility, since large interactive heatmaps require supercomputer-level memory resources to perform efficient, lag-free zooming and panning [36, 37, 38, 39, 40, 41, 45]; and unclear interpretation, since large heatmaps contain so much information that the standard recommended approach has been to preemptively subset the input data matrix into a smaller size [3].

Nevertheless, NGS-driven research studies often produce datasets on the order of 10^4^ rows (e.g., transcriptome studies such as the HTA 2.0 array [35] that have up to 400,000 rows, each representing individual exons). Likewise, single-cell RNA-seq studies often produce datasets ranging from several thousand to several hundred thousand cells [53, 54], posing significant computational challenges to efficient data visualization. Currently, interactively visualizing such big data is not possible using existing state-of-the-art methodologies, despite existing efforts in this direction [12, 13]. Unlocking the computational ability to visualize interactive heatmaps on such unprecedented size scales would allow researchers to investigate high-dimensional numerical data as a colored grid of cells that is easily zoomable to any desired resolution, thereby aiding the exploratory data analysis process.

With the advent of increasingly sophisticated interactive heatmap software and the rise of big data coupled with a growing community interest to examine it interactively, there has arisen an unmet and pressing need to address the computational limitations that hinder the production of large, interactive heatmaps. Examining such heatmaps would be valuable for visualizing the landscape of both global gene expression patterns as well as individual genes. Motivated to address these objectives, we propose an ultra fast and low memory user-friendly heatmap software suite capable of efficiently creating highly customizable static and interactive heatmaps in a web browser.

## Materials and Methods

shinyheatmap is hosted online as an R Shiny web server application. shinyheatmap may also be run locally from within R Studio, as shown here: https://github.com/Bohdan-Khomtchouk/shinyheatmap.shinyheatmap leverages the cumulative utility of R’s heatmaply [12], shiny [5], data.table [10], and gplots [50] libraries to create a cohesive web browser-based software experience requiring absolutely no programming experience from the user, or even the need to download R on a local computer. This kind of user-friendliness is geared towards the broader biological community, but will also appeal to the bioinformatics and computational biology communities. In contrast to most existing state-of-the-art heatmap software, shinyheatmap provides users with an extensive array of user-friendly hierarchical clustering methods, both in the form of multiple distance metrics as well as various linkage algorithms. This is especially useful for exploratory data analysis, particularly when the underlying data structure is unknown [31]. Since the choice of distance measure and linkage algorithm will directly influence the hierarchical clustering results, it is recommended to try different hierarchical clustering settings during analysis [31]. Agglomerative hierarchical clustering algorithms and their properties are described in detail at [27, 28, 44, 47, 15].

For the static heatmap generation, shinyheatmap employs the heatmap.2 function of the gplots library. For the interactive heatmap generation, shinyheatmap employs the heatmaply R package, which directly calls the plotly.js engine, in order to create fast, interactive heatmaps from large input datasets. The heatmaply R package is a descendent of the d3heatmap R package, which successfully creates advanced interactive heatmaps but is incapable of handling large inputs (e.g., 2000+ rows) due to memory considerations. As such, heatmaply constitutes a much-needed performance upgrade to d3heatmap, one that is made possible by the plotly R package [33], which itself relies on the sophisticated and complex plotly.js engine [24]. Therefore, it is the technical innovations of the plotly.js source code that make drawing extremely large heatmaps both a fast and efficient process. However, heatmaply also adds certain features not present in either the plotly.js engine nor the plotly R package, namely the ability to perform advanced hierarchical clustering and dendrogram-side zooming.

Despite these advantages, heatmaply is inadequate for plotting large datasets beyond a certain size limit, even with computationally expensive operations like hierarchical clustering disabled; for instance in certain cases, simple input matrices as small as 5000 × 5 may pose users with severe efficiency problems during heatmap rendering and zooming, even with no clustering present [13]. Due to this limitation, we developed a high performance web plug-in to shinyheatmap, called fastheatmap [11], which can rapidly plot interactive heatmaps of datasets as large as 10^5^ − 10^7^ rows within seconds directly in a web browser. Zooming in and out of such extremely large heatmaps is achievable in milliseconds, in contrast to d3heatmap or heatmaply, which takes minutes or even hours, if it is possible at all (due to memory limitations). This constitutes an unprecedented performance benchmark that dominantly positions shinyheatmap and its high performance computing server, fastheatmap, at the leading forefront of big data genomics heatmap visualization technology. In fact, to the best of our knowledge, the shinyheatmap/fastheatmap duo is the first big data software to appear on the biological heatmap visualization scene. All source code from the fastheatmap project is made publicly available at: https://github.com/Bohdan-Khomtchouk/fastheatmap.

## Results

To use shinyheatmap, input data must be in the form of a matrix of integer values. The value in the *i*-th row and the *j*-th column of the matrix denotes how many reads (or fragments, for paired-end RNA-seq) have been unambiguously assigned to gene *i* in sample *j* [21]. Analogously, for other types of assays, the rows of the matrix might correspond e.g., to binding regions (with ChIP-seq), species of bacteria (with metagenomic datasets), or peptide sequences (with quantitative mass spectrometry). For detailed usage considerations, shinyheatmap provides a convenient Instructions tab panel upon login.

Upon uploading the input dataset, both static and interactive heatmaps are automatically created, each in their own respective tab panel. The user can then proceed to customize the static heatmap through a suite of available parameter settings located in the sidebar panel (Figure 1). For example, hierarchical clustering, color schemes, scaling, color keys, trace, and font size can all be set to the specifications of the user. In addition, a download button is provided for users to save publication quality heatmap figures. Likewise, the user can customize the interactive heatmap through its own respective hoverable toolbar panel located at the upper right corner of the heatmap (Figure 2). This toolbar provides extensive download, zoom, pan, lasso and box select, autoscale, reset, and hover features for interacting with the heatmap. Users with large input datasets will be directed by shinyheatmap to its fastheatmap plug-in by way of a user-friendly message that automatically recognizes the dimensions of the input data matrix (Figure 3). Performance benchmarks indicate (Figure 4) that fastheatmap significantly outperforms the latest state-of-the-art interactive heatmap software by several orders of magnitude. All benchmarks were tested on a 64-bit Windows 10 Pro desktop machine with 16.0 GB of RAM and an Intel(R) Core(TM) i7-5820K CPU at 3.30 GHz.

**Figure 1.**
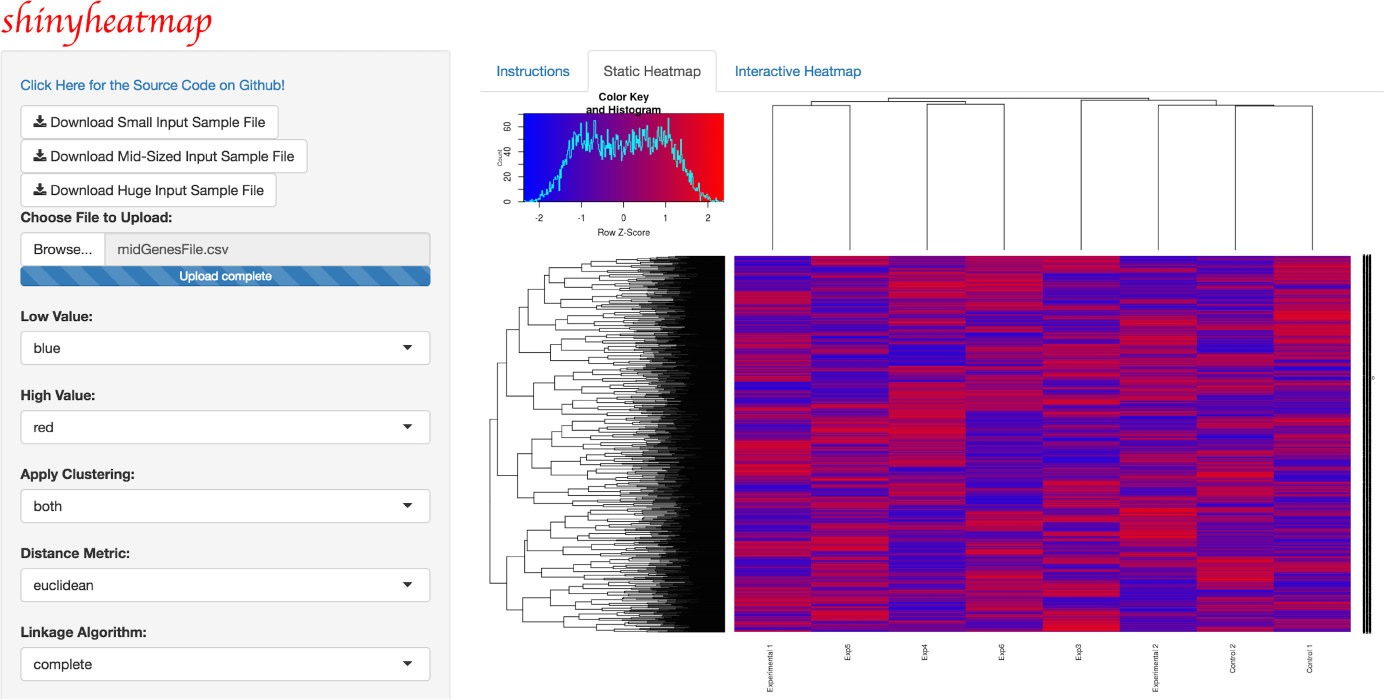
shinyheatmap static heatmap. shinyheatmap UI showcasing the visualization of a static heatmap generated from a large input dataset. Parameters such as hierarchical clustering (including options for distance metrics and linkage algorithms), color schemes, scaling, color keys, trace, and font size can all be set by the user. Progress bars appear during the heatmap rendering process to alert the user if any technical issues may arise. Sample input files of various sizes are provided as part of the web application, whose source code can be viewed on Github.

**Figure 2.**
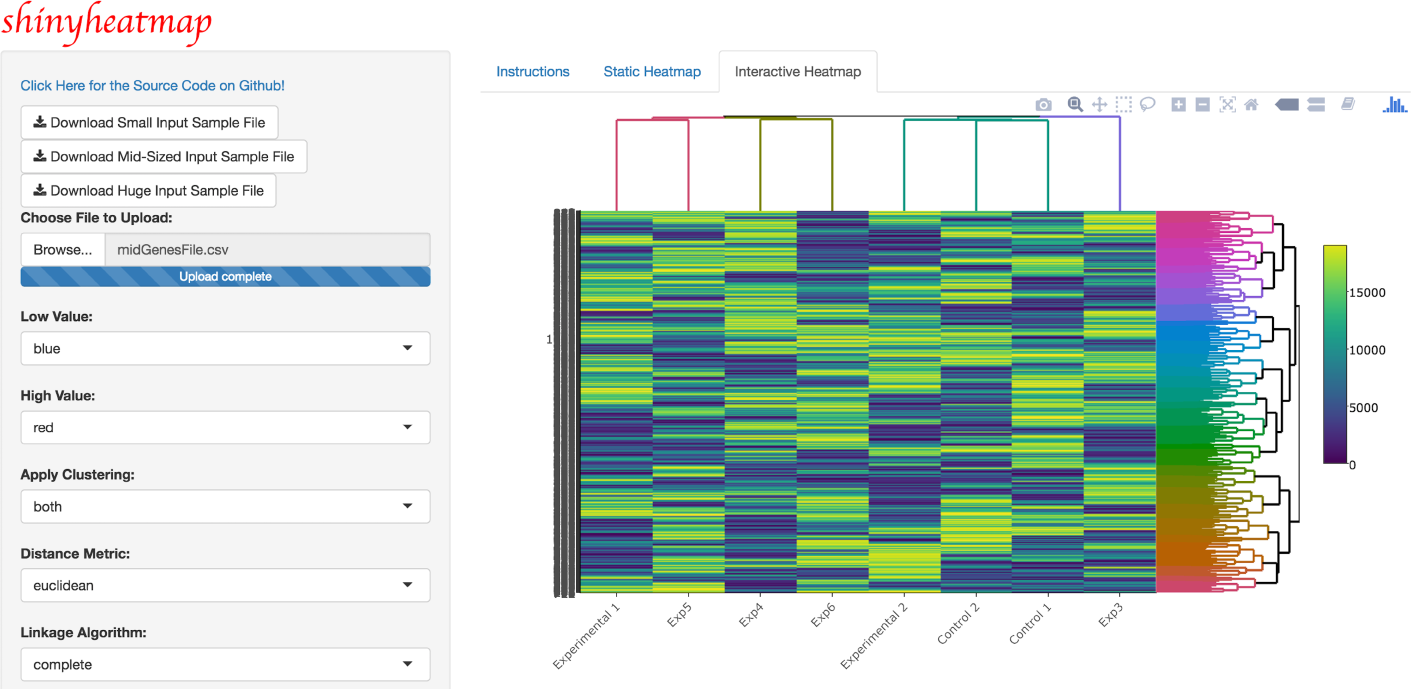
shinyheatmap interactive heatmap. shinyheatmap UI showcasing the visualization of an interactive heatmap generated from a large input dataset. An embedded panel that appears top right on-hover provides extensive download, zoom, pan, lasso and box select, autoscale, reset, and other features for interacting with the heatmap.

**Figure 3.**
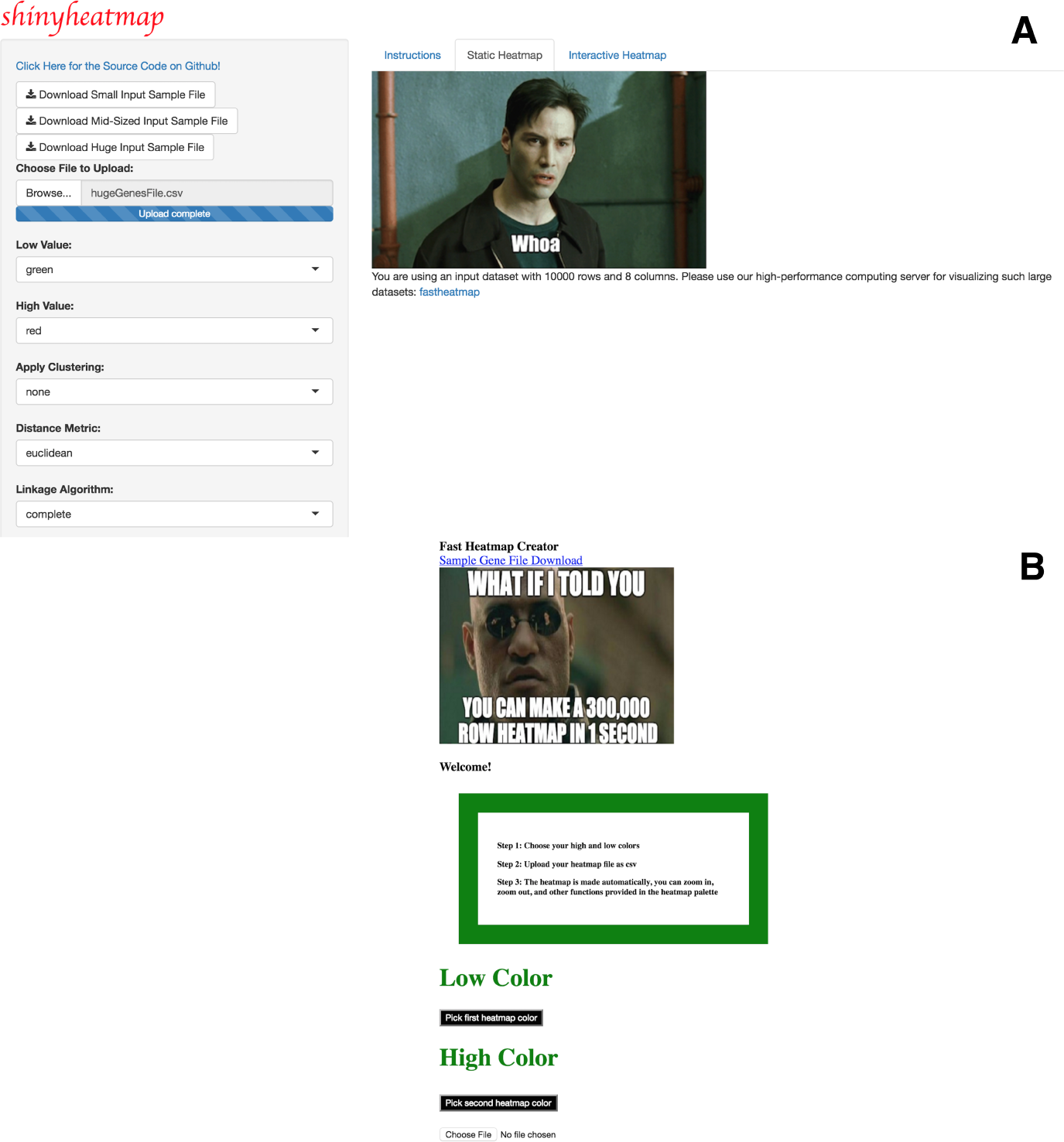
fastheatmap & shinyheatmap are linked together. A) shinyheatmap contains an auto-detector that detects the size of a user’s input matrix and, if the input matrix is too large, the user will be provided with a direct link to access shinyheatmap’s high performance computing server: fastheatmap. B) fastheatmap UI upon clicking on the URL link shown in Panel A.

**Figure 4.**
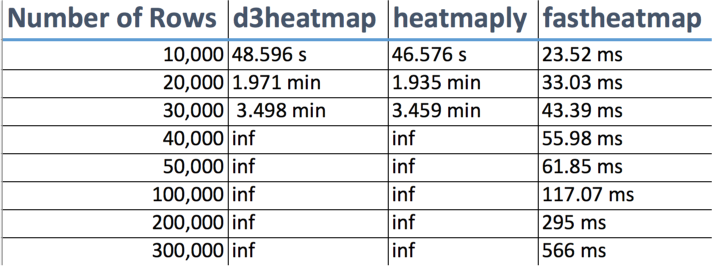
shinyheatmap performance benchmarks. shinyheatmap’s HPC plug-in, fastheatmap, performs > 100000 faster than other state-of-the-art interactive heatmap software. “Number of Rows” denotes the number of rows in the input file, “inf” (infinity) denotes a system crash due to memory overload, “s” denotes seconds, “min” denotes minutes, and “ms” denotes milliseconds.

## Conclusions

We provide access to a user-friendly web application designed to quickly and efficiently create static and interactive heatmaps within the R programming environment, without any prerequisite programming skills required of the user. Our software tool aims to enrich the genomic data exploration experience by providing a variety of customization options to investigate large input datasets.

## Declarations

### Author’s contributions

BBK conceived the study. BBK and JRH wrote the code. JRH benchmarked software. CW participated in the management of the source code and its coordination. BBK wrote the paper. All authors read and approved the final manuscript.

## Acknowledgements

BBK dedicates this work to the memory of his uncle, Taras Khomchuk. BBK wishes to acknowledge the financial support of the United States Department of Defense (DoD) through the National Defense Science and Engineering Graduate Fellowship (NDSEG) Program: this research was conducted with Government support under and awarded by DoD, Army Research Office (ARO), National Defense Science and Engineering Graduate (NDSEG) Fellowship, 32 CFR 168a. Relevant work in CW’s laboratory is currently funded by NIH grants DA035592 and AA023781.

## Ethics and consent to participate

This study does not involve humans, human data or animals.

## Consent to publish

Not applicable.

## Abbreviations used

HPC: high performance computing
PCA: principal component analysis
UI: user interface
URL: Uniform Resource Locator

## Availability of Data and Materials

All source code has been made publicly available on Github at:
https://github.com/Bohdan-Khomtchouk/shinyheatmap and https://github.com/Bohdan-Khomtchouk/fastheatmap.

